# Proteomics-based comparative mapping of the human brown and white adipocyte secretome reveals EPDR1 as a novel batokine

**DOI:** 10.1101/402867

**Authors:** Atul S. Deshmukh, Lone Peijs, Søren Nielsen, Rafael Bayarri-Olmos, Therese J. Larsen, Naja Z. Jespersen, Helle Hattel, Birgitte Holst, Peter Garred, Mads Tang-Christensen, Annika Sanfridson, Zachary Gerhart-Hines, Bente K. Pedersen, Matthias Mann, Camilla Scheele

## Abstract

Secreted proteins from adipose tissue play a role in metabolic cross-talk and homeostasis. We performed high sensitivity mass spectrometry-based proteomics on the cell media of in vitro differentiated, non-immortalized brown adipocytes derived from supraclavicular adipose of adult humans and white adipocytes derived from subcutaneous adipose of adult humans. We identified 471 potentially secreted proteins covering interesting protein categories such as hormones, growth factors, growth factor binding proteins, cytokines, extracellular matrix proteins, and proteins of the complement system, which were differentially regulated in brown and white adipocytes. A total of 101 proteins were exclusively quantified in brown adipocytes, among these ependymin-related protein 1 (EPDR1). Ablation of EPDR1 impaired the induction of thermogenic transcripts in response to norepinephrine in brown adipocytes, while EPDR1-treated mice increased their energy consumption, suggesting a role in brown fat commitment and activation. Our work reveals substantial differences between the secretomes of brown and white human adipocytes and identifies novel candidate batokines.

## Introduction

Adipose tissue is a major regulator of whole body energy homeostasis by communication with the brain and other organs (Stern et al., 2016). This cross talk is in part mediated by adipokines released from the adipocytes depending on the energy state. Well-established examples include leptin (LEP) and adiponectin (ADIPOQ), which are produced and secreted by white adipose tissue (WAT) (Ahima et al., 1996; Scherer et al., 1995). Since the discovery that adult humans have functionally competent brown adipose tissue (BAT) (Cypess et al., 2009; Saito et al., 2009; Van Marken Lichtenbelt et al., 2009; Virtanen et al., 2009; Zingaretti et al., 2009), there has been increasing interest in investigating BAT-derived adipokines, known as batokines (Villarroya et al., 2016). BAT produces heat in response to cold- or meal-induced sympathetic activation and increased BAT activity can counteract high-fat diet induced obesity in mice (Cannon and Nedergaard, 2004). In adult humans, the activity of BAT is negatively correlated with BMI (Saito et al., 2009; Van Marken Lichtenbelt et al., 2009) and several human in vivo studies suggest a functional role of BAT in whole-body metabolism (Scheele and Nielsen, 2017). Some of the identified batokines have been described as having hormonal function, enhancing BAT activity, improving glucose metabolism or mediating browning of white fat (Lee et al., 2014; Stanford et al., 2013; Svensson et al., 2016). Batokines could also be represented by growth factors acting in an autocrine or paracrine manner by regulating BAT differentiation (Villarroya et al., 2016). Mass spectrometry (MS)-based proteomics provides a powerful tool for the identification of novel secreted proteins from various cells including white adipocytes (Deshmukh et al., 2015; Meissner et al., 2013; Roca-Rivada et al., 2015); however, the human BAT secretome yet remains to be identified. Mapping factors secreted from human brown adipocytes would enhance our understanding of BAT-mediated metabolic organ cross-talk and might identify novel batokines which regulate BAT commitment and recruitment. The lack of representative human BAT cell models, as well as the challenges associated with measuring the secretome in cell culture media, have previously restricted these analyses. However, development of advanced secretomics technology as well as a non-immortalized human BAT cell model now make such experiments more feasible (Deshmukh et al., 2015; Jespersen et al., 2013; Meissner et al., 2013). In the current study, we investigate the human BAT and WAT secretomes using high-resolution MS-based proteomics to analyze cell culture media from brown and white human adipocytes. We mine the results for interesting novel candidates with potential hormonal activity, and perform follow up studies on EPDR1, a secreted modulator of energy homeostasis.

## RESULTS AND DISCUSSION

### The secretome of human brown and white adipocytes

Brown fat precursor cells were isolated from the supraclavicular adipose (termed brown throughout this manuscript) of adult humans (n=21), as previously reported (Jespersen et al., 2013). These non-immortalized cell cultures were differentiated in vitro and the five cultures (from different individuals) displaying the best differentiation capacity in terms of lipid accumulation and the highest induction of UCP1 expression in response to norepinephrine (NE) were included in the study. They were matched with white fat precursor cells derived from subcutaneous adipose (termed white throughout this manuscript) with equal differentiation capacity as assessed by visual estimation of lipid droplet accumulation and expression of the adipocyte differentiation marker fatty acid binding protein 4 (FABP4) (Figure 1A-C). Despite equal differentiation capacity and exposure to the same differentiation media formulation, we observed a higher uncoupling protein 1 (UCP1) expression in brown adipocytes both at baseline (Figure 1D) and following stimulation with NE (Figure 1E). These data align with previous studies from our laboratory demonstrating that isolated brown fat precursor cells maintain brown fat properties following in vitro differentiation (Gnad et al., 2014; Hiraike et al., 2017; Jespersen et al., 2013; Mottillo et al., 2016). Our cell model allows us to compare the secretomes between an unstimulated condition and the response to NE; a major activator of brown fat. The UCP1 expression underscores that these cell cultures are an appropriate model for the subsequent secretome studies. Thus we performed proteomics analysis of the serum-free cell culture media from these in vitro differentiated brown and white adipocytes. With high-resolution MS (Michalski et al., 2011) and automated computational analysis in MaxQuant (Cox and Mann, 2008) (workflow schematized in Figure 1F), we detected 1866 protein groups (protein entries that were distinguishable by MS of their identified peptides) in total in the conditioned cell culture media from brown and white adipocytes (with or without NE-stimulation) (Figure 1G, Table S1). In this set, computational analysis predicted 471 secreted proteins including 340 classical and 131 non-classically secreted proteins (Figure 1G, Table S1). We compared percentage of protein annotations before and after the computational analysis and as expected, we observed a decrease in the intracellular location category, whereas the categories related to protein secretion, such as glycoprotein, signal, and extracellular locations were increased (Figure 1H). The intracellular protein portion was still high after the bioinformatic analysis filtering for signal, secreted and extracellular protein annotations. However, these proteins had “multiple annotations” and besides the intracellular annotation, they either had a signal peptide (72.5%) or were annotated also as extracellular proteins (27.5%) (Figure 1H, Table S1). This is consistent with the idea that proteins may carry out different functions in different cellular locations (Huberts and van der Klei, 2010). Most of the secreted proteins were detected in cell culture media from both brown and white adipocytes (Figure 1G), but interestingly, vascular endothelial growth factor A (VEGFA) was only identified in the media from brown adipocytes while leptin (LEP) was exclusively identified in white adipocytes (Table S1). This provides proof of concept of our experimental workflow as LEP is one of the most studied white fat-derived hormones (Stern et al., 2016) while VEGFA is a well-established brown fat growth factor (Park et al., 2017; Shimizu et al., 2017).

**Figure 1.**
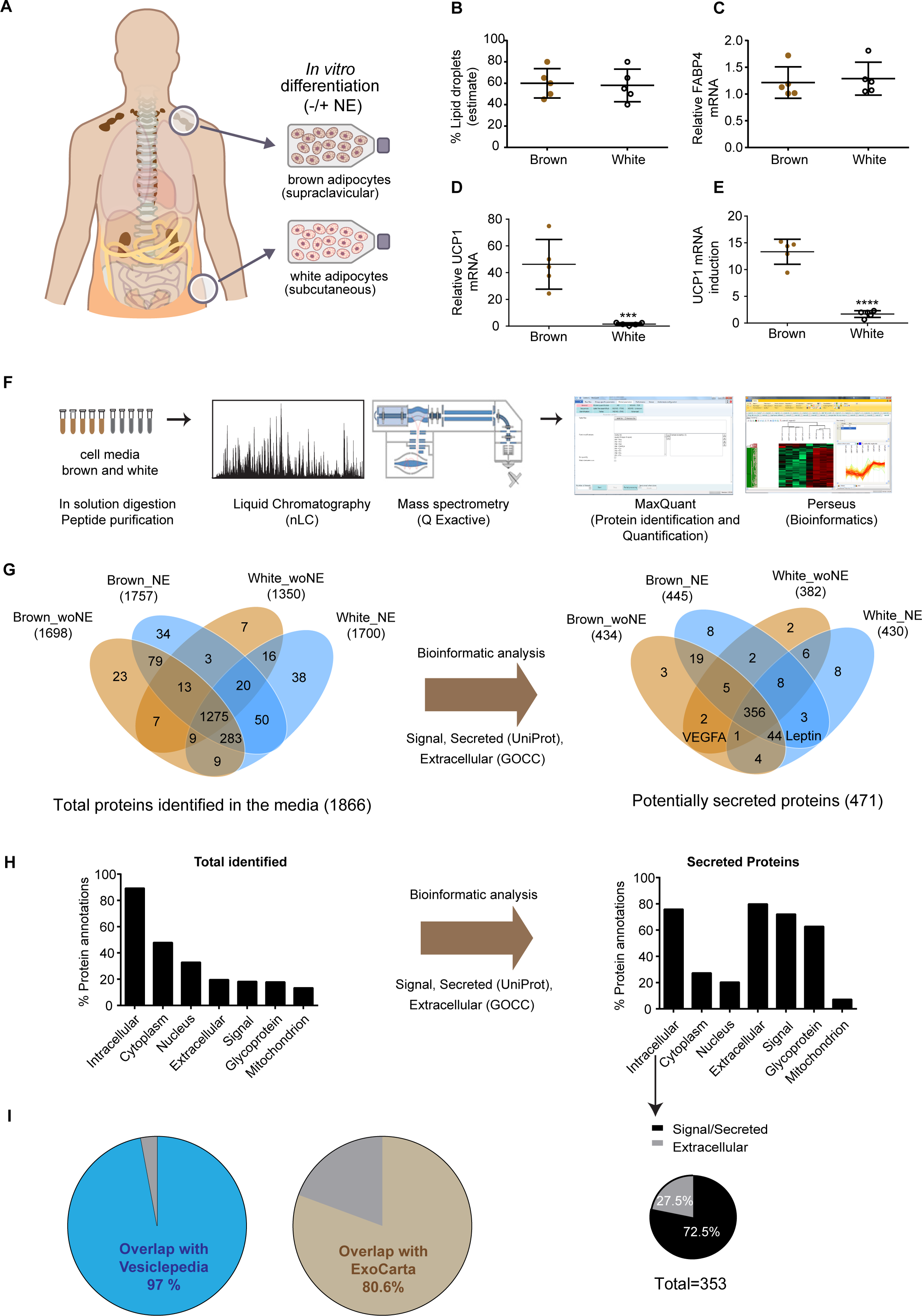
Model and proteomics workflow for generating the secretome of human brown and white adipocytes. **A)** Preadipocytes were isolated from the stromal vascular fraction of human supraclavicular adipose tissue biospies (brown adipose) and abdominal subcutaneous adipose tissue biopsies (white adipose), and were differentiated in vitro into lipid droplet containing adipocytes, using the same differentiation protocol for brown and white adipocytes. Mature adipocytes (brown adipocytes from n=5 human donors; white adipocytes from n=5 human donors) included in the study were characterized for **B)** Lipid droplet accumulation (estimated visually by phase contrast microscopy). **C)** FABP4 mRNA expression. **D)** UCP1 mRNA expression. **E)** UCP1 mRNA induction, calculated as fold change between unstimulated and norepinephrine (NE) stimulated adipocytes (4h stimulation). Data are mean +/-SEM; *P<0.05, **P<0.01, ***P<0.001. **F)** Proteomics workflow. **G)** Overlap of the total amount of identified proteins between the different conditions, before and after filtering for proteins predicted to be secreted. **H)** Categorization of identified proteins before and after filtering for proteins predicted to be secreted. **I)** Overlap between identified secreted proteins and databases of secreted proteins; Vesiclepedia and ExoCarta.

Whereas secretomes commonly are predicted based on the presence of a signal peptide suggesting secretion through the classical ER-Golgi pathway, hundreds of intracellular proteins are thought to be secreted through various non-classical pathways, perhaps including secretion from exosomes and microvesicles (Huberts and van der Klei, 2010; Nickel and Rabouille, 2009). We mapped our dataset with Vesiclepedia (Kalra et al., 2012) and ExoCarta (Keerthikumar et al., 2016), and found that almost all proteins overlapped with secreted proteins found in these two databases (Figure 1I). We performed live/dead staining and observed very limited cell death, further supporting the case that at least some of the proteins lacking signal peptides are truly secreted proteins.

### Comparative mapping of human brown versus white adipocyte secretomes

We compared the secretomes of the adipocytes isolated from the individual donors (n=5, supraclavicular and 5 subcutaneous) within the four groups (brown and white adipocytes with and without NE stimulation). We observed high correlations within the groups (median Pearson correlation for the brown adipocytes= 0.86, Figure 2A, Table S2). We applied label-free quantitation based on the MaxLFQ algorithm, which has proven robust and allows comparison of an arbitrary number of samples simultaneously (Cox et al., 2014). (Ong et al., 2002) (Roca-Rivada et al., 2015).

**Figure 2.**
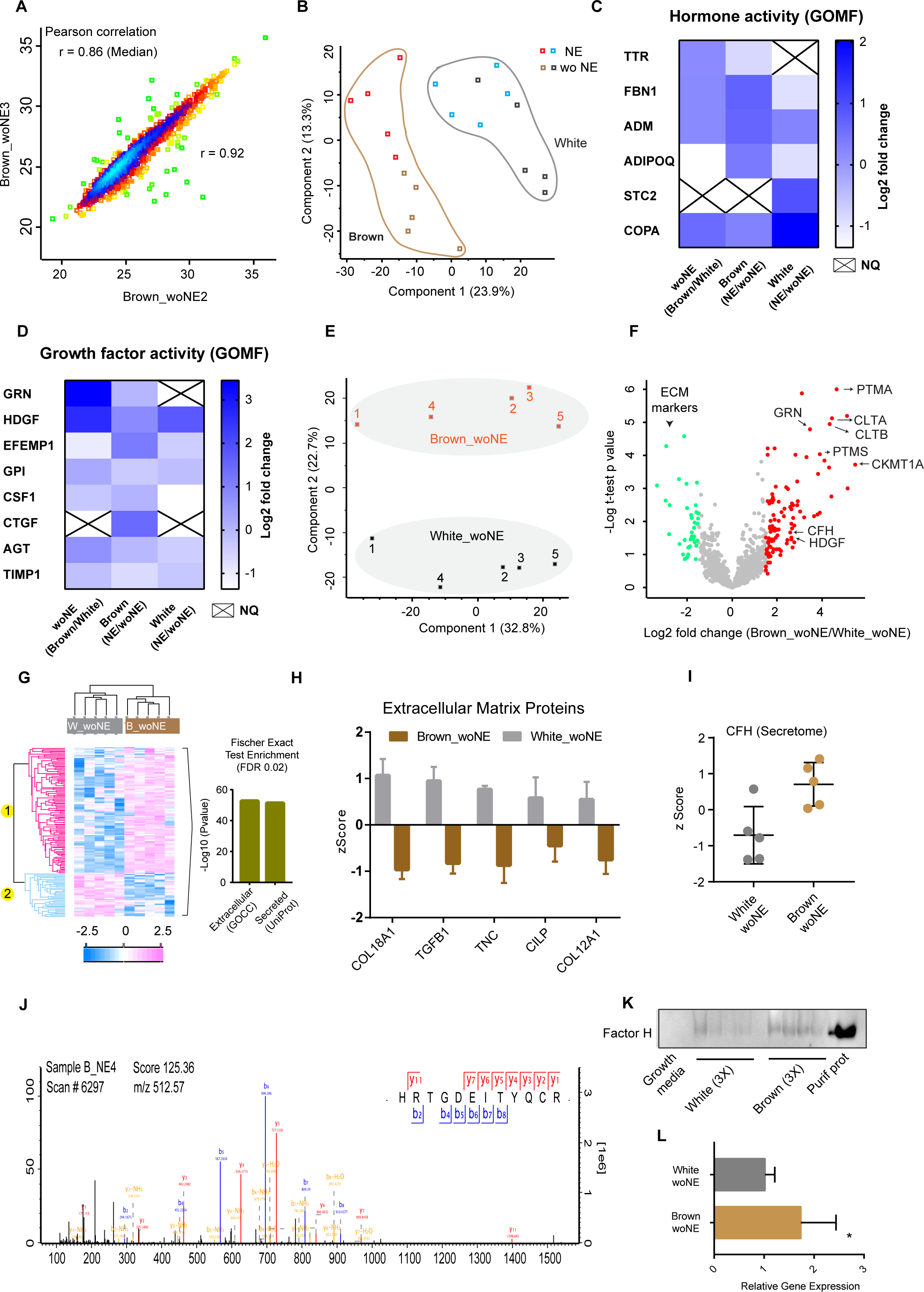
The quantified secretomes of human brown and white adipocytes. **A)** Representative Pearson correlation between two samples within the same group (brown adipocytes without NE stimulation). **B)** PCA plot including all proteins quantified in brown and white adipocytes with (NE) and without NE (wo NE) stimulation. **C)** Quantified proteins in the datasets with hormone activity, as annotated with the gene ontology molecular function (GOMF) term. NQ means not quantified. **D)** Quantified proteins in the datasets with growth factor activity, identified with GOMF term. **E)** PCA plot including all quantified proteins in unstimulated brown and white adipocytes. **F)** Two sample t-test was performed between all quantified proteins in unstimulated brown and white adipocytes and -Log t-test p-value were plotted against Log2 fold change. Green dots represent proteins that were higher abundant in cell media from white adipocytes and red dots represent proteins higher in brown adipocytes. **G)** Heatmap of all differentially regulated proteins between unstimulated brown and white adipocytes, as determined by the two sample t-test. **H)** ECM proteins, the main protein category higher expressed in white adipopcytes compared to brown adipocytes. **I)** Secreted amounts of Complement factor H (CFH) from brown and white adipocytes. **J)** Mass spectrometry trace of CFH. **K)** Western blot of CFH in brown and white adipocyte culture media from three different brown adipocyte cell strains and three different white adipocyte cell strains. **L)** CFH relative mRNA levels in mature lipid containing white and brown adipocytes measured by qPCR. Data are mean +/-SEM; *P<0.05, **P<0.01, ***P<0.001.

Human BAT and WAT adipose possess distinct metabolic properties (Scheele and Nielsen, 2017). To assess bioinformatically whether differences between brown and white adipocytes also were reflected in their secretomes, we performed principal component analysis (PCA). The analysis demonstrated a clear separation between the groups under stimulated and non-stimulated conditions (Figure 2B). Thus, the protein secretion patterns from brown and white adipocytes were sufficiently separated to classify them as distinct entities. Based on what has been reported on adipokines and batokines, we decided to systematically search our dataset for proteins that were annotated with gene ontologies (GO) for growth factor activity or hormone activity. This identified six hormones (Figure 2C) and eight growth factors (Figure 2D). We calculated the fold change for these proteins between unstimulated brown and white adipocytes (woNE Brown/White); unstimulated and NE-stimulated brown adipocytes (Brown NE/woNE) and between unstimulated and NE-stimulated white adipocytes (White NE/woNE). Among the hormones were the well-described adipokine, adiponectin (ADIPOQ; log2 fold changes: -1.35; 0.41; -0.92) (Scherer et al., 1995; Stern et al., 2016). Interestingly, we also quantified Fibrillin-1 (FBN1; log2 fold changes: 0.20; 0.72; -0.93), the precursor protein for the newly discovered adipose hormone, asprosin (Duerrschmid et al., 2017; Romere et al., 2016). Proteins not previously described as adipokines or batokines included: transthyretin (TTR; log2 fold changes: 0.18; -0.84; N/A), a binding protein for thyroid hormone which in turn has been shown to induce lipid oxidation in BAT (Seoane-collazo et al., 2017); adrenomedullin (ADM; log2 fold changes: 0.19; 0.67; 0.27), which is cleaved into adrenomedullin (AM) and proadrenomedullin (PAMP), peptides with vasodilative properties; stanniocalcin-2 (STC2; log2 fold changes: N/A; N/A; 0.95), regulating calcium and phosphate homeostasis and coatomer subunit alpha (COPA; log2 fold changes: 0.60; 0.30; 2.03), which is cleaved into two peptides; xenin and proxenin. Among the growth factors, we observed that granulins (GRN; log2 fold changes: 3.45; 0.001; N/A) and hepatoma-derived growth factor (HDGF; fold changes: 2.64; 0.82; 1.75) were more abundant in the brown adipocyte media compared to the white adipocyte media and thus are candidate batokines. Granulins are cleaved into nine chains and have been described as autocrine growth factors important for wound-healing (He et al., 2003). Granulins have not previously been described as batokines. However, one of the nine chains, acrogranin, also known as progranulin, was identified as a WAT-derived adipokine mediating high-fat diet-induced insulin resistance in mice (Matsubara et al., 2007). HDGF promotes mitogenic activity through DNA-binding mediated transcriptional repression (Yang and Everett, 2007), but has also been detected in the extracellular region (Nüße et al., 2017).

Taken together, we here quantified both established adipokines and novel candidate batokines and adipokines. To gain insights into the differences between secretomes of brown and white adipocytes, we performed a two-sample t-test (False Discovery Rate (FDR) < 0.05). When comparing NE-induced secretomes in both brown and white adipocytes, we surprisingly observed many ribosomal proteins, or proteins involved in translational processes (Table S4) following NE stimulation. We decided to focus on the proteins secreted from brown and white adipocytes without norepinephrine stimulation in our search for novel batokines. A PCA plot including all proteins quantified under non-stimulated conditions revealed a clear separation between brown and white adipocytes (Figure 2E). Two-sample t-test returned 143 differentially regulated proteins of which 106 were more abundant in cell media from brown adipocytes while 37 were more abundant in cell media from white adipocytes (Figure 2F, 2G, Table S4). Fischer exact test (FDR: 0.02) on signficantly different protein in the background of total quantified proteins (1113) returned only two categories; ‘Extracellular (GOCC)’ and ‘Secreted (UniProt Keyword)’. This analysis suggests that the majority of differentially regulated proteins in brown and white adipocyte cell culture media are secreted proteins (Figure 2G). Several white adipocyte-selective secreted proteins were ECM-associated, including Transforming growth factor beta 1 (TGFB1); which has been associated with diabetes risk (Kim et al., 2013) and tenascin (TNC); an ECM glycoprotein with proinflammatory effects, which is highly expressed in WAT of obese patients and in mice models of obesity (Kim et al., 2013). We observed a more than two-fold higher secretion of TNC and COL18A1 in cell media from white adipocytes compared to brown adipocytes (Figure 2G, 2H, Table S4). Cartilage intermediate layer protein 1 (CILP) and Collagen alpha-1(XII) chain (COL12A1) are additional extracellular matrix proteins accumulating in the white adipocyte cell media. It would be interesting to further study the biological role of these secreted collagens and other ECM associated proteins in the WAT/BAT context.

In the cell media of brown adipocyte cell media there were highly abundant proteins covering diverse functions. Parathymosin (PTMS) and prothymosin alpha (PTMA) were more than four-fold higher in cell media from brown adipocytes compared with white adipocytes (Figure 2F). These relatively small proteins (11-12 kDa) were quantified with more than five unique peptides in cell media from brown adipocytes (Table S4) and have been shown to be involved in proliferation and regulation of immune function (Hannappel and Huff, 2003; Samara et al., 2017). The biological role of PTMS and PTMA in brown fat has not been explored. Although these proteins are not annotated as secreted, it is possible that they are secreted via exosomes or microvesicles from brown adipocytes. Interestingly, the Clathrin-related proteins CLTA and CLTB, coating protein of intracellular vesicles (Pearse, 1976), were also among the proteins with higher abundance in cell media from brown adipocytes compared to white adipocytes. This suggests that brown adipocytes make use of alternative secretion pathways such as vesicular trafficking more frequently than white adipocytes. Further, we observed that mitochondrial creatine kinase U-type (CKMT1) was more abundant in the media from brown adipocytes. This protein is involved in BAT mitochondrial energy metabolism and has recently gained interest as a human BAT-selective protein (Kazak et al., 2015; Müller et al., 2016).

Among of the proteins that were secreted in higher amounts from the brown adipocytes was a negative regulator of the innate immune system, complement factor H (CFH) (Figure 2F, 2I-L). CFH inhibits the alternative pathway of the complement system by binding to C3b and enhancing the degradation of the alternative C3 convertase (C3bBb). In addition, CFH acts as a cofactor to complement factor I (CFI), a central inhibitor of all three complement system pathways, i.e. the classical, lectin and alternative pathways (Zipfel and Skerka, 2009). Therefore, our MS data of the adipocyte media suggest that brown adipocytes might have higher anti-inflammatory capacity compared to white adipocytes. To further investigate this hypothesis, we performed western blots on cell culture media from the brown and white adipocytes. We reproduced the findings from the MS analysis as we found that CFH was present in the cell culture media and generated a stronger signal in the media from brown adipocytes compared to white adipocytes (Figure 2K). In concordance, mRNA expression of CFH was higher in brown adipocytes as opposed to white adipocytes (Figure 2L).

### A distinct secretome of brown adipocytes

Next, we investigated secreted proteins that were specifically quantified in the cell culture media from either brown or white adipocytes. The majority of proteins (262) were quantified in cell culture media from both brown and white adipocytes; but a subset of the proteins were exclusively quantified in cell culture media from brown adipocytes (101) or white adipocytes (37) (Figure 3A, Table S5). The brown adipocyte-specific proteins are interest and relevant to investigate as batokine candidates, because they may mediate inter-organ cross talk. One protein that specifically caught our interest was mammalian ependymin-related protein 1 (EPDR1) (Figure 3B-C).

**Figure 3.**
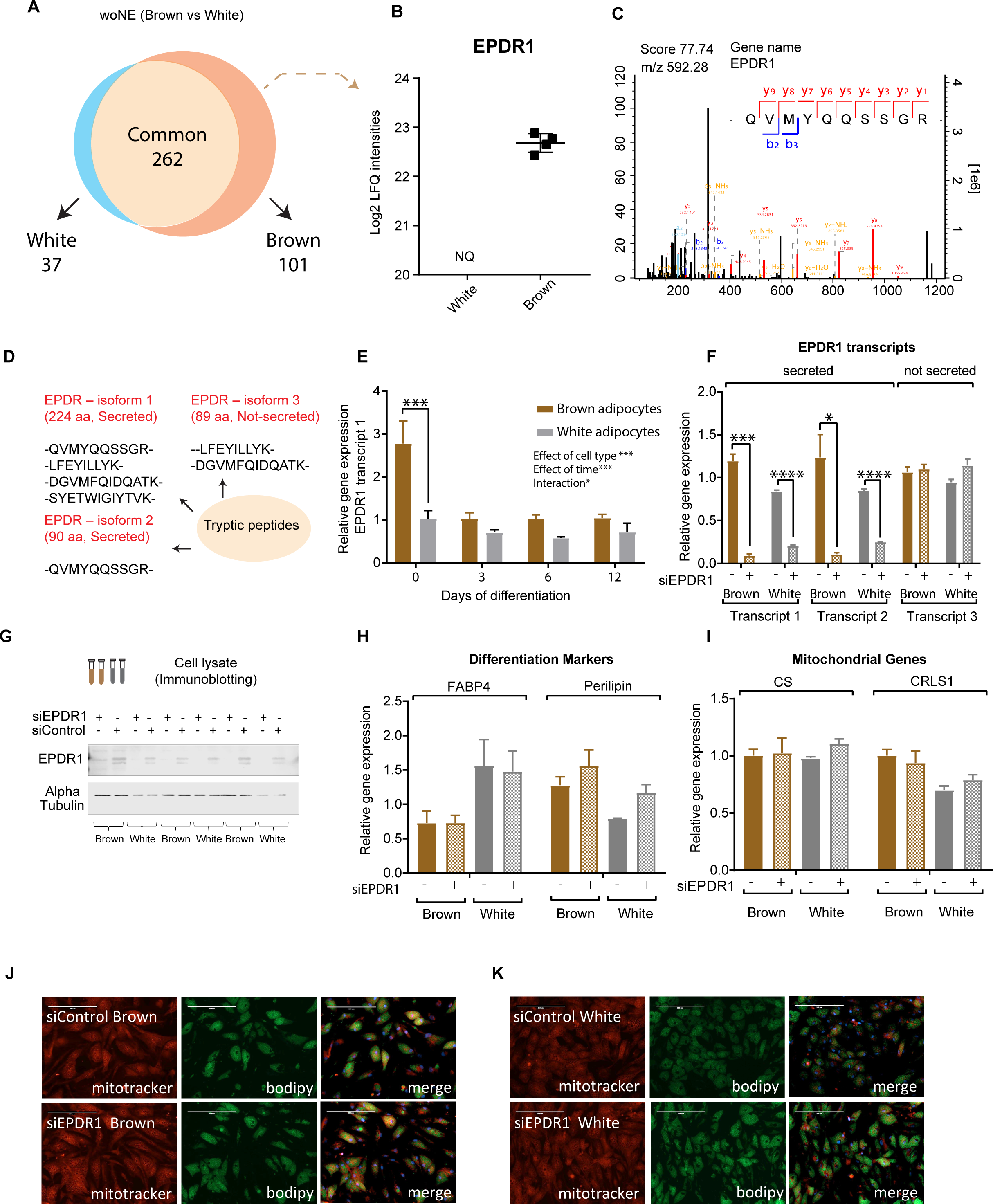
Characterization of EPDR1, a novel batokine candidate. **A)** Venn diagram illustrating proteins annotated as secreted based on gene ontology cellular component (GOCC) enrichment analysis for extracellular annotations in combination with Uniprot keyword annotations for predicted signal sequences or secretion on all detected proteins. **B)** Protein quantification of EPDR1 in cell media from white and brown adipocytes. NQ means not quantified. **C)** MS trace of peptides mapping to EPDR1. **D)** Peptides mapping to EPDR1 depicting different transcript variants. **E)** Gene expression of EPDR1 transcript variant 1 during differentiation of brown or white adipocytes. **F)** Gene expression of EPDR1 transcript variants in siEPDR1 or siControl transfected adipocytes. **G)** Western Blot for EPDR1 and α-Tubulin on EPDR1 siRNA (siEPDR1) or Non-targeting siRNA (siControl) transfected adipocytes. **H)** Gene expression on fat differentiation markers FABP4 and PLIN1 on siEPDR1 or siControl transfected adipocytes. **I)** Gene expression on general mitochondrial markers CS and CRLS1 on siEPDR1 or siControl transfected adipocytes. **J)** Mitochondrial (mitotracker) and lipid droplet (bodipy) staining in siEPDR1 or siControl transfected brown adipocytes. **K)** Mitochondrial (mitotracker) and lipid droplet (bodipy) staining in siEPDR1- or siControl-transfected white adipocytes. (Data are presented as mean of means ± SEM (n=3 independent experiments with one cell strain), p-value *<0.5**<0.01*** <0.001****<0.0001). Statistical significance was determined by two-way ANOVA with matching values and Sidaks post hoc test for multiple comparisons.

EPDR1 was previously identified in a deep sequencing and bioinformatic study of PR/SET domain 16 (PRDM16)-induced beiging in mice (Svensson et al., 2016). EPDR1 is a so far functionally undescribed protein, but it has been shown to be differentially expressed during adipocyte differentiation in a proteomic investigation of murine adipocytes (Ye et al., 2011). Very recently, EPDR1 gene expression was shown to be downregulated in subcutaneous human fat biopsies during short- and long-term weight loss and displayed opposite expression patterns in acquired obesity (Bollepalli et al., 2017). The *EPDR1* gene encodes three transcript variants, translated to isoforms two of which include an N-terminal signal peptide for secretion. We detected peptides unique to the secreted isoform 1, confirming its presence in our samples. Peptides mapping to isoform 2 (secreted) and isoform 3 (not secreted) were also detected, but these peptides also mapped to isoform 1, thus their presence in our samples could neither be confirmed nor excluded (Figure 3D and Figure S1). To assess the expression levels of the transcript variants encoding the isoforms we designed transcript-specific qPCR assays and found that the expression of transcript variant 1 (encoding isoform 1) was higher than the two other transcript variants both with and without NE stimulation in the differentiated brown adipocytes (data not shown). During differentiation, EPDR1 transcript variant 1 was most highly expressed at day 0 of the 12-day differentiation program in brown adipocytes, and at this time point, the difference in EPDR1 expression between brown and white adipocytes was highest (Figure 3E). To begin to investigate the functional role of EPDR1, we performed an siRNA mediated knockdown of EPDR1 in brown and white adipocyte cell strains, targeting only the transcripts containing a signal peptide for secretion. This approach led to a sustained decrease in EPDR1 as confirmed by transcript-specific qPCR and Western blot 12 days after transfection in fully differentiated adipocytes (Figure 3F-G). There were no changes in gene expression of the two differentiation markers FABP4 and perilipin 1 (PLIN1) (Figure 3H) or in the mitochondrial markers citrate synthase (CS) and cardiolipin synthase (CRLS1) between the control and EPDR1 knockdown cells (Figure 3I). Finally, we did not observe any visual changes in lipid droplet accumulation or mitochondrial content (Figure 3J). This indicates that the reduction of EPDR1 did not affect the brown or white adipogenesis in terms of lipid accumulation and mitochondrial biogenesis.

### EPDR1 is important for brown fat commitment

We next assessed the effect of EPDR1 knockdown at day 0 of differentiation, on NE-response in fully differentiated brown and white adipocytes. Knockdown of EPDR1 at the onset of differentiation, had no effect on the early brown fat co-transcription factor PRDM16, but interestingly inhibited the NE-induction of the thermogenic transcriptional program including UCP1, PPARG coactivator 1 alpha (PGC1-α) and iodothyronine deiodinase 2 (DIO2) in brown adipocytes (Figure 4A-D). These data indicated that EPDR1 acts downstream of PRDM16, a master regulator of brown adipogenesis (Seale et al., 2011). This is consistent with the previous finding that PRDM16 overexpression in inguinal fat increases EPDR1 secretion in mice (Svensson et al., 2016). In white adipocytes, expression of UCP1, PGC-1α and DIO2 were not altered by EPDR1 knockdown. To investigate the functional effect of the observed phenotype, we assessed the effect of EPDR1 knockdown on the NE-induced proton leak. Basal respiration was not changed, but NE-induced proton leak was reduced following knockdown of EPDR1 in brown adipocytes (Figure 4E-F). These findings indicated that EPDR1 is important for brown fat commitment during early differentiation of brown preadipocytes, as ablation in early differentiation resulted in a functionally deficient brown fat phenotype.

**Figure 4.**
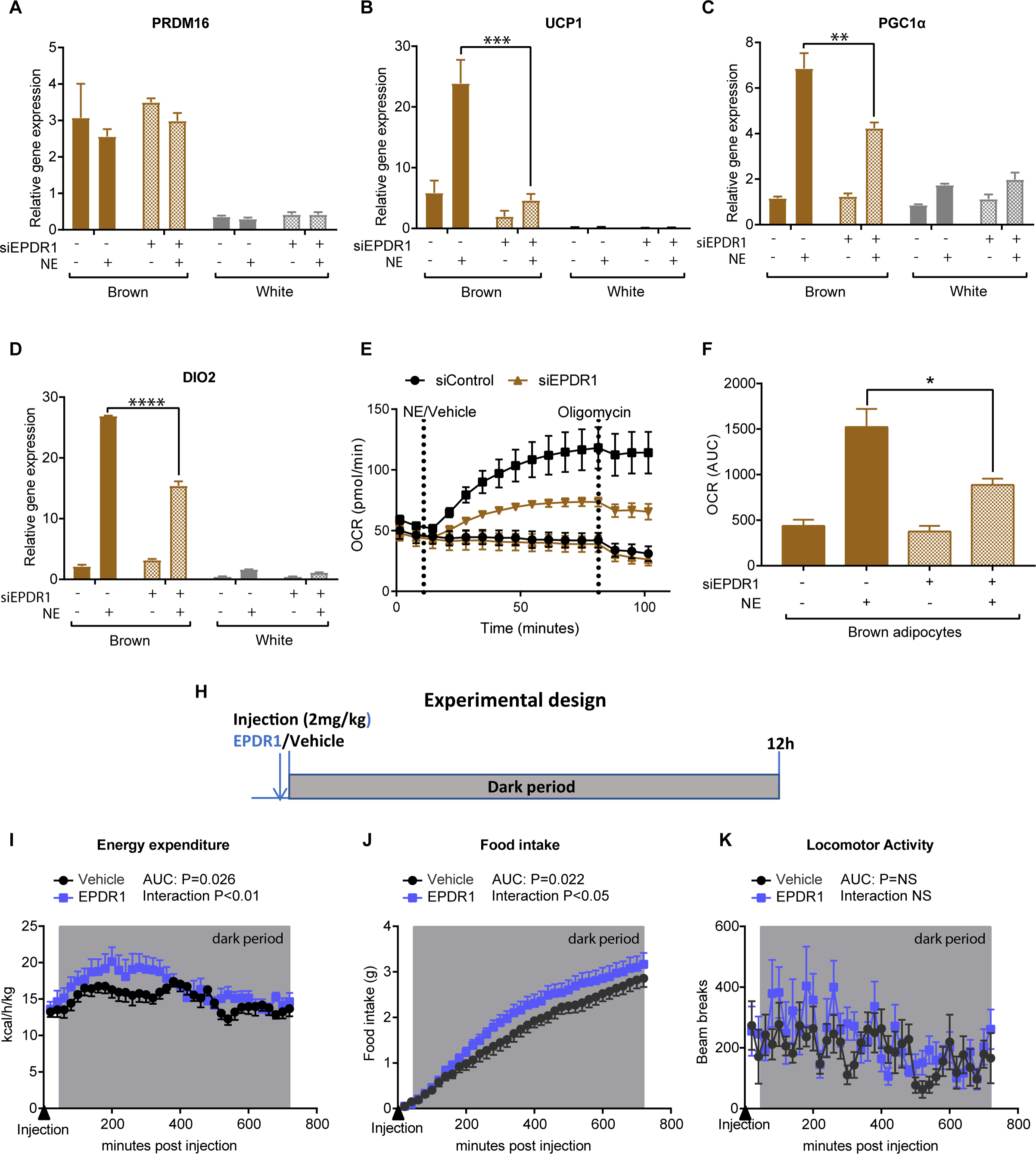
The role of EPDR1 in brown fat differentiation and activity. **A)** Gene expression of PRDM16 in mature adipocytes stimulated with vehicle or NE (4h) and transfected with EPDR1 siRNA (siEPDR1) or Non-targeting siRNA (siControl) at day 0. **B)** Gene expression of UCP1 in mature adipocytes stimulated with vehicle or NE and transfected with siEPDR1 or siControl. **C)** Gene expression of PGC-1α in mature adipocytes stimulated with vehicle or NE and transfected with siEPDR1 or siControl. **D)** Gene expression of DIO2 in mature adipocytes stimulated with vehicle or NE and transfected with siEPDR1 or siControl. (data are presented as mean of means ± SEM (n=3 independent experiments), p-value *<0.5**<0.01****<0.0001). Statistical significance was determined by two-way ANOVA with matching values and Sidaks post hoc test for multiple comparisons. **E)** Oxygen consumption rate in brown adipocytes transfected with siEPDR1 (black squares and circles) or siControl (brown triangles) (n=3 independent experiments with one cell strain). **F)** Area under the curve AUC on oxygen consumption rates in brown adipocytes transfected with siEPDR1 or siControl following oligomycin treatment. **G)** Experimental design for injection of EPDR1 in C57Bl/6 mice (n=8). **H)** Calculated energy expenditure following EPDR1 (blue) or vehicle (black) injection during the dark period. **I)** Food intake following EPDR1 (blue) or vehicle (black) injection during the dark period. **J)** Locomotor activity following EPDR1 (blue)or vehicle (black) injection during the dark period. Statistical significance was determined by two-way ANOVA with repeated measures and the p-value for interaction is reported. Area under the curve (AUC) was calculated and the p-value is reported in the figure.

Finally, we addressed whether EPDR1 also had acute effects on whole body metabolism. We injected 14-week-old C57BL/6 mice, which had been acclimated at thermoneutrality (30^0^C) for 18 days prior to the experiment to ensure minimal physiological adrenergic stimulation (Jacobsson et al., 1994), with EPDR1 protein at 2 mg/kg (Figure 4G).. This environment is more representative of human conditions and allowed us to assess whether EPDR1 had any effect on the metabolism. The injection was given just prior to the dark period where brown fat activity is naturally increased by murine circadian rhythm (Gerhart-Hines et al., 2013). We recorded whole-body energy expenditure throughout the dark period using indirect calorimetry. Subcutaneous injection of EPDR1 protein resulted in an increase in energy expenditure during a 12-hour period as compared with vehicle-injected control mice (Figure 4H). This increase in energy expenditure was followed by a subsequent increase in food intake (Figure 4I), exemplifying the fine-tuned synergy between energy expenditure and energy intake (Contreras et al., 2017, 2014; Sutton et al., 2014). There was no difference in locomotor activity (Figure 4J) during the 12-hour period, and respiratory exchange ratio (RER) was unchanged, excluding physical activity as a driving factor of the increased energy expenditure. Hence, our data indicate that increased circulating EPDR1 augments whole body energy expenditure independently of physical activity. Instead, the increase in energy expenditure might be related to an acutely enhanced thermogenic activity, which would suggest a dual role of EPDR1 in brown fat commitment and activity.

In conclusion, we here provide the first comprehensive analysis of the human brown fat secretome compared to the white fat secretome and their response to NE. Our results reinforce that brown and white adipocytes have distinct secretory profiles and metabolic functions. We provide a large number of novel candidate batokines. Among several interesting candidates, we focused on a novel human batokine, EPDR1, which is selectively secreted from brown adipocytes, although it transcript is present in both white and brown adipocytes. We demonstrate that EPDR1 regulates the thermogenic gene program and function in brown adipocytes and that subcutaneous injection of EPDR1 increases whole body energy expenditure in mice. Thus, the human BAT secretome represents a promising source of novel metabolic regulators.

## Methods

### Human Primary Adipocyte Cultures

Brown fat precursor cells were isolated from the supraclavicular region of a cohort of 20 adult humans also including tissue biopsies which have been previously described along with a subset of the cell cultures (Jespersen et al., 2013). White fat precursor cells were obtained from the subcutaneous abdominal region of another five adult humans. All subjects provided written informed consent. The Scientific-Ethics Committees of the Capital Region and Copenhagen and Frederiksberg Municipalities Denmark approved the study protocols, journal numbers H-A-2009-020, H-A-2008-081, and (KF) 01-141/04, respectively, and the studies were performed in accordance with the Helsinki declaration. Cells were plated and maintained in DMEM F12 (Thermo Fisher Scientific) containing 10% FBS, 1% Penicillin/Streptomycin and 1nM FGF1 (Immunotools). Two days post confluence (designated day 0) the cells were induced to differentiate by replacing the medium with DMEM/F12 with 10 µg/ml Transferrin, 2 nM T3, 100 nM Insulin, 100 nM Dexamethasone, 200 nM Rosiglitazone and 540 µM 3-Isobutyl-1-methylxanthine (IBMX). At day 3, the medium was replaced with the same medium composition, with the exemption of IBMX. The cells were refed with new medium at day 6 and day 9 with the exemption of Rosiglitazone. At day 12, cells were considered fully differentiated mature adipocytes. Secretome analysis was performed on supraclavicular (n=5) and subcutaneous (n=5) cells. For secretome analysis, fully differentiated cells were washed with DMEM/F12 and subsequently incubated for 2 h in DMEM/F12 containing 1% penicillin-streptomycin. Serum-starved cells were stimulated with 10 µM norepinephrine (NE) (Sigma-Aldrich) for 4 h. Cell culture media (2 ml) was collected for secretome analysis while cells were harvested for RNA analysis. To address whether cell viability was affected by NE treatment, cells were stained according to the manufacturer’s protocol, applying a LIVE/DEAD fluorescent assay (Thermo Scientific) visualized using an EVOS FL fluorescent microscope (Thermo Scientific).

### siRNA mediated knockdown of EPDR1

At day 0 of the differentiation program, adipocytes were transfected with 22.2 nM siRNA targeting EPDR1 or a non-targeting pool (Dharmacon) using Lipofectamine^®^ RNAiMAX Reagent (Thermo Fisher Scientific) according to the manufacturer’s instructions. The cells were then cultured as described in the previous section.

### RNA Isolation and Quantitative Real-Time PCR

Total RNA from human adipocytes was isolated with TRIzol (Thermo Fisher Scientific), according to the manufacturer’s recommendations. RNA concentrations were measured using Nanodrop 1000, and 250 ng of RNA was used as input for subsequent cDNA synthesis with High Capacity cDNA Synthesis Kit (Thermo Fisher Scientific). Primer sequences can be found in Table S6 Target mRNA was normalized to PPIA and calculated using the comparative delta-delta-Ct method.

### Oxygen consumption measurements

Mitochondrial respiration rates were assessed using the XFe96 Extracellular Flux Analyzer (Agilent Technologies). Cells were plated at confluence (7000cells/well) and transfected and differentiated as described above. The cell medium was replaced with Seahorse Base medium without phenol red, supplemented with 1 µM L-Glutamine, 2 µM Na_2_PO_4_ and 25 mM Glucose pH 7.4 1 h prior to respiration measurements. Oxygen consumptions rates were followed under basal conditions, NE (1 µM) injection and finally oligomycin injection (20 µM).

### Lipid and mitochondrial staining

Mature adipocytes were incubated with DMEM/F12 containing 0.2µM MitoTracker Red CMXRos (Thermo Fisher Scientific) for 20 min. Cells were then washed 3 times with PBS and fixed with 4% formaldehyde (Sigma-Aldrich) for 15 min. Fixed cells were washed again 3 times with PBS, and lipid were stained with 0.5 mM Bodipy (Thermo Fisher Scientific) for 20 min. Subsequently nuclei were stained with NucBlue Fixed ReadyProbe Reagent (Thermo Fisher Scientific) for 7 min. Cells were washed 3 times with PBS and visualized with EVOS FL imaging system (Thermo Fisher Scientific).

### Secretome Analysis by Mass Spectrometry

We used high-resolution mass spectrometry (MS)-based approach to detect a highly complex secreted protein mixture from human primary brown and white adipose cell cultures. In our workflow, serum free cell supernatants were trypsin digested, and the resulting peptide mixtures were directly analyzed in a single-run LC-MS format. We performed liquid chromatography with 2 h gradient and analyzed peptides on bench top quadrupole-Orbitrap instrument with very high sequencing speed and high mass accuracy in MS and MS/MS modes (Michalski et al., 2011). Label-free quantification of the MS data is performed in the MaxQuant environment while bioinformatics analysis was done with Perseus software (Tyanova et al., 2016).

#### Secretome digestion

Secretome analysis of the conditioned media from brown and white adipocytes was performed as described before (Deshmukh et al., 2015) with slight modifications. Proteins in conditioned media were denatured with 2 M urea in 10 mM HEPES pH 10 by ultrasonication on ice and acetone precipitated (overnight). Protein pellets were washed with 80% acetone and suspended in urea (6 M) and thiourea (2 M) buffer (pH 8). Proteins were reduced with 10 mM dithiothreitol for 40 min followed by alkylation with 55 mM iodoacetamide for 40 min in the dark. Proteins were digested with 0.5 µg LysC (Wako) for 3 h and digested with 0.5 µg trypsin for 16 h at room temperature. The digestion was stopped with 0.5% trifluoroacetic acid, 2% acetonitrile. Peptides were desalted on reversed phase C18 StageTips. The peptides were eluted using 20 µl of 60% acetonitrile in 0.5% acetic acid and concentrated in a SpeedVac. Concentrated peptides were acidified with 2% acetonitrile, 0.1% trifluoroacetic acid in 0.1% formic acid.

#### LC MS/MS analysis

The peptides were analyzed using LC-MS instrumentation consisting of an Easy nanoflow UHPLC (Thermo Fischer Scientific) coupled via a nanoelectrospray ion source (Thermo Fischer Scientific) to a Q Exactive mass spectrometer (Thermo Fischer Scientific) (Michalski 2011). Peptides were separated on a 50 cm column with 75 µm inner diameter packed in-house with ReproSil-Pur C18-aq 1.9 µm resin (Dr. Maisch). Peptides were loaded in buffer containing 0.5% formic acid and eluted with a 160 min linear gradient with buffer containing 80% acetonitrile and 0.5% formic acid (v/v) at 250 nL/min. Chromatography and column oven (Sonation GmbH) temperature were controlled and monitored in real time using SprayQC (Scheltema and Mann, 2012). Mass spectra were acquired using a data-dependent Top8 method, with the automatic switch between MS and MS/MS. Mass spectra were acquired in the Orbitrap analyzer with a mass range of 300-1650 m/z and 70 000 resolution at m/z 200. HCD peptide fragments were acquired with a normalized collision energy of 25. The maximum ion injection times for the survey scan and the MS/MS scans were 20 and 220 ms, and the ion target values were set to 3e6 and 1e5, respectively. Data were acquired using Xcalibur software.

### Computational MS data analysis

The raw files for secretome and cellular proteome were analyzed in the MaxQuant environment (Tyanova et al., 2016). The initial maximum allowed mass deviation was set to 6 ppm for monoisotopic precursor ions and 20 ppm for MS/MS peaks. Enzyme specificity was set to trypsin, defined as C-terminal to arginine and lysine excluding proline, and a maximum of two missed cleavages was allowed. A minimal peptide length of six amino acids was required. Carbamidomethylcysteine was set as a fixed modification, while N-terminal acetylation and methionine oxidation were set as variable modification. The spectra were searched by the Andromeda search engine against the human UniProt sequence database with 248 common contaminants and concatenated with the reversed versions of all sequences. The false discovery rate (FDR) was set to 1% for peptide and protein identifications. The peptide identifications across different LC-MS runs were matched by enabling the ‘match between runs’ feature in MaxQuant with a retention time window of 30 s. If the identified peptides were shared between two or more proteins, these were combined and reported in protein group. Contaminants and reverse identifications were removed from further data analysis. Protein quantification was based on the Max LFQ algorithm integrated into the MaxQuant software (Cox et al., 2014).

Bioinformatics analysis was performed with the Perseus software (Tyanova et al., 2016) (http://www.perseus-framework.org). Categorical annotation was supplied in the form of KEGG pathways, Keywords (UniProt), and Gene Ontology (GO) (biological process (BP), molecular function (MF) and cellular component (CC)). All annotations were extracted from the UniProt database. To define the secretome of brown and white adipocytes, we applied previously described computational workflow on all proteins identified in the media (Deshmukh et al., 2015). Briefly, signal peptide-containing proteins were categorized as ‘classical’ secreted proteins while protein annotated to ‘extracellular location’ (GOCC) or ‘secreted’ (UniProt, Keywords) were classified as ‘non-classical’ secreted proteins. While comparing proteins identified in the cell media from brown or white adipocytes with Vesiclepedia (Kalra et al., 2012) and ExoCarta databases(Keerthikumar et al., 2016), we included only evidence at the protein level.

The global comparative analysis was performed on LFQ intensities. We used very stringent criteria for MaxLFQ-based quantification (min ratio count 2 in MaxQuant). Moreover, we included only those proteins which were quantified at least 2 times in at least one group (i.e. Brown_woNE vs White_woNE) Due to the randomness of peptide sampling in shotgun proteomics, the quantification of several proteins is missing for some samples. The data was imputed to fill missing abundance values by drawing random numbers from a Gaussian distribution. These parameters have been tuned in order to simulate the distribution of low abundant proteins best. To investigate differences between brown and white adipocyte secretomes, we compared brown and white adipocyte media proteome under non-stimulated (woNE) conditions while the effect of NE-stimulation was investigated by comparing stimulated and non-stimulated conditions (Brown_woNE vs Brown_NE/ White_woNE vs White_NE). These comparisons were made using Two sample t-test in Perseus with FDR 0.05. Proteins which were regulated by 1.5-fold (Log2) were considered as significantly different proteins (Table S4). Two-sample t-test (volcano plot), hierarchical clustering and annotation enrichment were based on label-free quantitation of the samples. Hierarchical clustering of significantly different proteins was performed after Z-score normalization. We then performed Fisher exact test on significantly different protein (background total quantified proteins), testing for enrichment or depletion of any annotation term in the cluster compared to the whole matrix.

### Western Blot Analysis on cell media

For complement factor H, we validated proteomics data using western blot analysis on conditional media. The conditioned media was concentrated (3 different Brown adipocyte cell strains and 3 different White adipocyte cell strains) through two centrifugation rounds using an Amicon Ultra-4 3K and 10 K filter devices (Merk Millipore, USA). Purified proteins (FB Hycult HC2129, FH CompTech A137, FI CompTech A138) were used as positive control. Growth media was used as negative control. Precision Plus Protein Blue (Bio-Rad, USA) was used as molecular weight (MW) marker. The CHF antibody was produced in-house by Prof. Peter Garred’s research group using mouse hybridomas.

### Western Blot Analysis on cell lysate

SiRNA mediated knockdown of EPDR1 was validated using western blot analysis on cell protein lysate. In short, the mature differentiated cells were washed twice with ice cold PBS and the lysed in lysis buffer containing 20 mM tris-HCl, 150 mM NaCl, 1 mM EDTA, 1 mM EGTA, 1% Triton-X, 2.5 mM Na_4_P_2_O_7_, 1 mM β-glycerophosphate, 1mM Na_3_VO_4_, Complete^(™)^ Mini, Protease Inhibitor Cocktail. Lysates were centrifuged for 15 min at 13.000 g at 4^°^ C. Protein concentration was determined using the Bio-Rad Protein (Bio-Rad, California, USA). 3.6 µg of protein lysate was loaded on Bis-Tris SDS-page gels and subject to electrophoresis, using iBright Prestained Protein ladder (Thermo Fisher Scientific) to determine MW of detected bands. Proteins were transferred to PVDF membranes by semi-dry transfer for 7 min with Pierce Power blot cassette. Membranes were blocked in FSG and incubated for 16 h with anti-EPDR1 (Thermo Fisher Scientific) or anti-α-Tubulin (Sigma-Aldrich). Bands were detected with IRDye secondary antibodies (LI-COR) and visualized using the Odyssey^®^ Fc Imaging System (LI-COR)

### Animal studies

All animal studies were approved by the Animal Experimentation Inspectorate of the Danish Ministry of Justice, no 2014-15-0201-00181. Twelve-week old male C57BL/6NRJ mice (Janvier Labs, France) were used for the animal studies. The animals were single housed and maintained at a 12-h light-dark cycle (03:00/15:00) in temperature (30°C–32°C) and humidity (50–60%) controlled rooms, with free access to standard chow (Altromin 1314F, pellets) and tap water. For metabolic measurements, the TSE phenomaster (TSE systems GmbH, Bad Homburg, Germany) were used. The system utilizes indirect calorimetry to estimate energy expenditure from the mouse oxygen (VO2) consumption and carbondioxide (VCO2) production. All animals were acclimatized in habituation cages one week before they were placed in the TSE system. Here they were acclimatized for another 4 days prior to the initiation of the injection experiments and measurements. Mice were randomized according in two groups of eight mice taking their weight into account (mean body weight 25.8 g and 25.6 g for vehicle and EPDR1 treated group respectively). Recombinant EPDR1 protein was produced by Novo Nordisk A/S, Måløv, Denmark. EPDR1 (2 mg/kg in PBS) or PBS as vehicle was administered subcutaneously 20 min prior to the dark cycle. Metabolic measurements were followed for a total of 43 hours, and the initial 12-hour dark period was used to assess treatment effects.

### Statistical Analysis

Statistical analysis of proteomics data was performed in Perseus software (Tyanova Nature methods 2016). Statistical analysis for assessing differences in mRNA levels was performed using GraphPad Prism software. Data were presented as mean, and error bars represent SEM. Nonparametric Mann-Whitney tests were used for comparisons between the groups. Sample size and significance levels for each analysis are stated in figure legends. A p-value of 0.05 was considered significant.

## DATA ACCESSION NUMBER

The mass spectrometry proteomics data have been deposited to the ProteomeXchange Consortium via the PRIDE partner repository with the dataset identifier PXD008541. Reviewer account details: **Username:**, **Password:** WDqMl1Aa

## SUPPLEMENTAL INFORMATION

**Supplemental Table S1**: Total proteins identified in the brown and white adipocytes media under non-stimulated and stimulated conditions. The table also includes the list of predicted classical and non-classical secreted proteins (+).

**Supplemental Table S2:** Pearson correlation *r* between the replicates.

**Supplemental Table S3:**Median abundance (log2 LFQ intensities) of proteins with hormone or growth factor activities.

**Supplemental Table S4:** Quantitative differences between brown and white adipocytes secretomes under stimulated and non-stimulated conditions. The table also includes the fold change and whether proteins are up or down-regulated.

**Supplemental Table S5:** Median abundance (log2 LFQ intensities) of secreted proteins quantified in brown and white cell media. The list also includes proteins exclusive to BAT and WAT.

**Supplemental Table S6:** Primer sequences

## Acknowledgements

We thank Jacob Steen Petersen and Birgitte Andersen from Novo Nordisk A/S for valuable scientific discussion and for producing the EPDR1 protein used in the animal studies. The Centre for Physical Activity Research (CFAS) is supported by TrygFonden (grants ID 101390 and ID 20045). During the study period, the Center of Inflammation and Metabolism (CIM), Rigshospitalet, was supported by a grant from the Danish National Research Foundation (DNRF55). CIM/CFAS is a member of DD2, the Danish Center for Strategic Research in Type 2 Diabetes (the Danish Council for Strategic Research, grant nos. 09-067009 and 09-075724). The Novo Nordisk Foundation Center for Basic Metabolic Research (http://www.metabol.ku.dk) is supported by an unconditional grant from the Novo Nordisk Foundation to University of Copenhagen. The study was further funded by a shared research grant from Novo Nordisk A/S to CS and MM.

## Author contributions

CS and MM supervised the study. CS, ASD, MM, LP, SN, BKP, ZGH, AS, MT, PG, BH: hypothesis generation, conceptual design, data analysis, and manuscript preparation. LP, ASD, SN, TJL, RBO, NZJ, HH: conducted experiments and data analysis. All authors edited and approved the final manuscript.

## Figure legends

**Figure S1. Protein sequences of EPDR isoforms (source – UniProt).** The protein sequences of EPDR1 isoforms are displayed and the tryptic peptides detected by MS are mapped to the sequences.

